# Case-only analysis of gene-environment interactions using polygenic risk scores

**DOI:** 10.1101/555300

**Authors:** Allison Meisner, Prosenjit Kundu, Nilanjan Chatterjee

## Abstract

Investigations of gene (*G*)-environment (*E*) interactions have led to limited findings to date, possibly due to weak effects of individual genetic variants. Polygenic risk scores (PRS), which capture the genetic susceptibility associated with a set of variants, can be a powerful tool for detecting global patterns of interaction. Motivated by the case-only method for evaluating interactions with a single variant, we propose a case-only method for the analysis of interactions with a PRS in case-control studies. Assuming the PRS and *E* are independent, we show how a linear regression of the PRS on *E* in a sample of cases can be used to efficiently estimate the interaction parameter. Furthermore, if an estimate of the mean of the PRS in the underlying population is available, the proposed method can estimate the PRS main effect. Extensions allow for PRS-*E* dependence due to associations between variants in the PRS and *E*. Simulation studies indicate the proposed method offers appreciable gains in efficiency over logistic regression and can recover much of the efficiency of a cohort study. As an illustration, we apply the proposed method to investigate interactions between a PRS and epidemiologic factors on breast cancer risk in the UK Biobank study.

---

Many diseases have complex etiologies, including both environmental and genetic factors and their interactions (1,2). Understanding gene (*G*)-environment (*E*) interactions enhances our ability to model risk and thereby identify high-risk subgroups and could provide insights into biological mechanisms of disease (2–5). Consequently, the study of *G-E* interactions has been a focus in many areas of research (5), including psychiatry (6), pulmonology (7), and oncology (8–11). Here, we consider binary outcomes and focus on interactions as departures of joint effects from multiplicative relative risks. When the data come from a case-control study, the standard approach is to test for interaction using logistic regression; when the outcome is rare, the resulting odds ratio estimates will approximate the relative risks.

The development of powerful analytic methods for investigating *G-E* interactions in case-control studies has been an active area of research for several decades (12). One of the most important contributions in this line of research was the development of case-only method for estimating *G-E* interactions for categorical *G* and *E* (13). Under G-E independence in the underlying population and a rare disease assumption, the interaction odds ratio can be estimated by regressing *E* on *G* (or vice versa) in a sample of cases (13). This approach can yield large gains in efficiency relative to logistic regression, which does not incorporate the independence assumption (13,14). Motivated by the potential for gains in efficiency by exploiting *G-E* independence, a number of novel methods were subsequently developed that extended the case-only method, including methods based on the retrospective likelihood (14, 15), data-adaptive shrinkage methods (16,17), and two-stage hypothesis testing procedures for genome-wide scanning of interactions (2,18). In spite of these developments, genome-wide association studies (GWAS) have thus far discovered few *G-E* interactions (4,5). It is likely that because individual genetic variants have very weak effects, the power to identify *G-E* interactions at the level of individual variants is low even with large sample sizes and relatively efficient methods.

Findings from GWAS indicate many complex diseases are highly polygenic, the result of the joint action of a large number of variants (19). Thus, polygenic risk scores (PRS), which quantify the genetic risk from a set of single-nucleotide polymorphisms (SNPs) (19), offer an attractive alternative to single-variant analysis of *G-E* interactions. PRS aggregate existing knowledge, removing the need to test for interactions between *E* and the individual genetic markers that comprise the PRS. Furthermore, PRS are expected to have more variability in the underlying population than single variants, which may lead to increased power for detecting interactions (3,4,20). Using PRS to investigate *G-E* interactions may be most fruitful if many SNPs have similar types of interactions; in the absence of such similarity, the interactions between the individual SNPs and *E* may become diluted when the SNPs are combined into the PRS. In settings where the goal is to develop a parsimonious risk prediction model including the joint effects of genetic and non-genetic factors, using PRS to evaluate interactions is particularly compelling (e.g., (21–24)).

In this paper, we demonstrate that exploiting independence between a PRS and E leads to a case-only method for studying the interaction between the PRS and *E* and that this approach offers gains in efficiency over logistic regression. Specifically, we show this interaction can be evaluated using a simple linear regression of the PRS on *E* in a sample of cases. We then show how the main effect of the PRS can be estimated and how the method can be extended to the setting where the PRS and *E* are correlated due to associations between *E* and some of the variants in the PRS. We use breast cancer data from the UK Biobank, a large cohort study, to illustrate application of the proposed method.

## METHODS

Throughout, we consider a binary outcome *D*, where *D* is either 0 or 1, with 1 indicating disease. Parameter estimates are denoted by a circumflex (or “hat”).

### PRS definition

A PRS is typically defined as a weighted sum of risk alleles for a collection of SNPs, Σ*_i_w_i_G_i_*, where *w_i_* is the weight for the *i*^th^ SNP and *G_i_* indicates the number of copies of the risk allele for the *i*^th^ SNP (0, 1, or 2) (19). The weight *w_i_* is typically an estimate of the association between the *i*^th^ SNP and the outcome, such as an estimated log odds ratio. We focus on the scenario where the weights are given, e.g., are based on summary statistics from large GWAS, and need not be estimated.

### *G-E* independence

Suppose the true population risk model for a *D* conditional on the PRS for *D, S_D_*, and *E* is

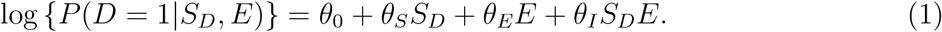

Furthermore, suppose

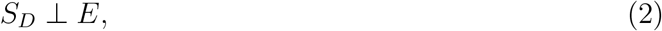

that is, *S_D_* and *E* are independent in the population. The parameter *θ_I_* characterizes the interaction between *S_D_* and *E* on the log relative risk scale. Since *S_D_* is a weighted sum of a number of SNPs, we assume the central limit theorem holds and the distribution of *S_D_* can be characterized by a normal distribution with mean *μ* and variance *σ*^2^. Under independence of *S_D_* and *E*, we have (*S_D_|E*) ~ *S_D_ ~ N*(*μ,σ*^2^). Using this result and the model given in equation 1, we demonstrate (Web Appendix A, available at https://academic.oup.com/aje) that (*S_D_|E,D* = 1) ~ *N*(*μ + σ*^2^(*θ_S_ + θ_I_E*),*σ*^2^); in other words, the conditional distribution of *S_D_* among cases is normal, where the mean is a linear function of *E*.

Thus, in practice, we can evaluate the interaction between *S_D_* and *E* by fitting a linear regression to a sample of cases:

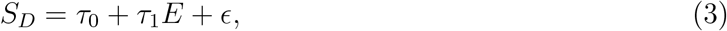

where *ϵ* ~ *N*(0,*σ*^2^). This provides estimates of *τ*_0_ = *μ* + *σ*^2^*θ_S_* and *τ*_1_ = *σ*^2^*θ_I_*. Scaling 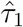 by 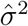 yields 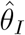. An estimate of *σ*^2^ could be obtained internally, e.g., based on the residual standard error estimate from the model in equation 3. Alternatively, as *σ*^2^ is also the marginal variance of *S_D_*, an external sample or, if the disease is rare, a sample of controls could be used to estimate *σ*^2^. The estimated standard error for 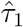 must also be scaled by 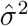 to provide proper inference for 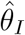 (Web Appendix B). When the population model is the log-risk model given in equation 1, if the outcome is rare for all levels of *E*, the parameter characterizing the interaction between *S_D_* and *E* on the log odds ratio scale is approximately equal to *θ_I_* (Web Appendix C).

#### Estimating the main effect of S_D_

In the linear regression model defined in equation 3, *τ*_0_ = *μ* + *σ*^2^*θ_S_*. Since *μ* is the marginal mean of *S_D_*, if an external sample is available or if the disease is rare and a sample of controls is available, this sample could be used to estimate *μ*. This, combined with 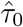 and 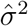, can be used to provide 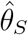. Since a finite sample is used to estimate *μ*, the estimated standard error of 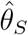 must be adjusted to account for variability in 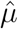, though the need for adjustment diminishes as the size of the sample used to estimate *μ* grows (Web Appendix D). As with the interaction parameter, if the outcome is rare for all levels of *E*, the parameter characterizing the main effect of *S_D_* on the log odds ratio scale is approximately equal to *θ_S_*.

### Accounting for *G-E* dependence for heritable *E*

In some settings, correlation between *S_D_* and *E* may arise as some endogenous environmental factors may have a heritable component. For example, body mass index (BMI) is a strong risk factor for many chronic diseases and recent GWAS have identified hundreds of SNPs associated with BMI (25). Thus, in exploring interactions between BMI and *S_D_* on the risk of *D*, one needs to be cautious about the potential for *G-E* correlation due to variants in *S_D_* that are also associated with BMI. In the following, we propose a way of correcting for such *G-E* dependence by utilizing a PRS for *E*.

Let us suppose, in addition to *S_D_*, we can calculate *S_E_*, a PRS for *E*. The SNP weights used in *S_E_* could be obtained from a large GWAS for *E*. We will assume that conditional on *S_E_, S_D_* is independent of *E*, which implies *S_E_* captures all of the association of the variants in *S_D_* with *E*. We further assume that conditional on *S_D_* and *E, D* and *S_E_* are independent, i.e., *S_E_* provides no independent information about disease risk given *S_D_* and *E*.

We assume the multivariate central limit theorem holds and the joint distribution of (*S_D_, S_E_*) can be characterized by a bivariate normal distribution with means *μ* and *δ*, respectively, variances *σ*^2^ and *γ*^2^, respectively, and correlation λ. Then (*S_D_|S_E_*) ~ *N*(*ϕ*_0_ + *ϕ*_1_*S_E_, ψ*^2^), where *ϕ*_0_ = *μ* − *σ*λ*δγ*^−1^, *ϕ*_1_ = *σ*λ*γ*^−1^, and *ψ*^2^ = *σ*^2^(1 − λ^2^). Using arguments similar to those used above in the setting of *G-E* independence, it can be shown (*S_D_|E, S_E_, D* =1) ~ *N*((*ϕ*_0_ + *ϕ*_1_*S_E_*) + *ψ*^2^(*θ_S_* + *θ_I_E*),*ψ*^2^). Thus, we can study *θ_I_* by fitting a linear regression to a sample of cases:

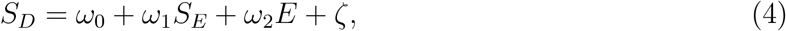

where *ζ* ~ *N*(0,*ψ*^2^).

This provides estimates of *ω*_0_ = *ϕ*_0_ + *ψ*^2^*θ_S_, ω*_1_ = *ϕ*_1_ and *ω*_2_ = *ψ*^2^*θ_I_*. By scaling 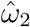 by 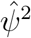 (e.g., based on the residual standard error estimate from the model in equation 4), we can obtain 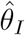. The standard error estimate for 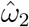 must be similarly scaled by 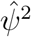 in order to provide proper inference for 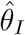. If 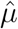 and 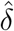 are also available, *θ_S_* can be estimated using 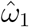, since *ω*_0_ = *μ* − *δω*_1_ + *ψ*^2^*θ_S_*.

### UK Biobank data

To illustrate application of our method, we used data on breast cancer from the UK Biobank (26,27), a large cohort study that enrolled approximately 500,000 individuals aged 37-74 years and is following them to observe health outcomes. The UK Biobank data included 264,232 women. We sought to explore *G-E* interactions on risk of post-menopausal breast cancer, defined as breast cancer after age 50 years. We defined our cohort as unrelated women of white British ancestry who did not have pre-menopausal breast cancer, were aged at least 50 years at study entry, and did not have breast cancer before entry. This yielded a cohort with 116,030 individuals, of which 2,339 were incident cases. The UK Biobank was approved by the North West Multi-centre Research Ethics Committee. This research was conducted under UK Biobank Resource application 17712.

Previous work evaluated a 77-SNP breast cancer PRS in a case-control study with 28,239 cases and 30,445 controls (24). The PRS was categorized into deciles and interactions between the PRS and several variables, including alcohol intake, height, and BMI, were considered. Significant interactions were found for alcohol intake and height, but clear dose-response patterns were not seen. In other words, the estimated associations between the environmental variables and breast cancer risk did not uniformly increase or decrease with increasing PRS decile.

We generated a PRS based on a recent breast cancer GWAS conducted by Michailidou et al. (28) This study identified 161 SNPs that achieved genome-wide significance (*P* value ≤ 5×10^−8^); of these, 151 SNPs were available in the UK Biobank data. Thus, we constructed a 151-SNP breast cancer PRS based on the summary statistics provided by Michailidou et al. and the genotype data available from the UK Biobank. We considered interactions between the PRS and alcohol intake derived from self-reported consumption (defined as g/day), BMI, and height. All variables were modeled as continuous variables. We compared three analyses: the case-only method with all 2,339 incident cases, an analysis of a mock case-control study with all 2,339 incident cases and 2,339 randomly sampled controls, and an analysis of the full cohort of 116,030 women, including the 2,339 incident cases. For the case-control and cohort analyses, we fit a logistic regression model and a Poisson regression model with robust variance estimation (29), respectively. The case-control and cohort analyses included adjustment for age at study entry. All analyses were conducted using R, version 3.5.1 (R Foundation for Statistical Computing, Vienna, Austria).

Assuming the true population risk model is similar to the model in equation 1, to the extent breast cancer risk is low for all levels of a given *E* during the follow-up period of the cohort (mean follow-up time = 5.7 years), all three methods will estimate approximately the same interaction parameter, i.e., the interaction on the relative risk scale. Otherwise, the case-control approach will estimate the interaction on the odds ratio scale, a distinct parameter. Given the low prevalence of breast cancer in the population and relatively modest observed associations between the variables considered here (height, BMI, and alcohol consumption) and breast cancer risk (30,31), it is anticipated that all three approaches will estimate approximately the same interaction parameter.

### Simulations

We conducted simulations to investigate the ability of the proposed case-only method to estimate and provide inference for the parameters of interest. Throughout, we compared our method to a case-control study and a cohort study; all three approaches (case-only, case-control, and cohort) included the same number of cases. We fit a logistic regression model for the case-control approach and a Poisson model with robust variance estimation for the cohort approach. In all simulations, the true population risk model was the log-risk model in equation 1 and the parameter values used were such that the outcome was rare for all levels of E; thus, all three approaches (approximately) estimate the interaction parameter and the *S_D_* main effect parameter on the log relative risk scale.

For all simulations, we first generated a population of 10^6^ observations. From this population, a sample of *n* cases and *n* controls were drawn; the *n* cases were used by the case-only method, while the *n* cases and *n* controls were used in the case-control approach. For the cohort approach, a sample of size *n/p*, where p was the proportion of cases in the population, was drawn; this yielded a sample with *n* cases and the same disease prevalence as in the population, on average. Throughout, *n* was 1000, 2000, or 5000. All simulations were repeated 1000 times. All *P* values (used to estimate the type I error rate) were two-sided and the nominal type I error rate was 0.05.

#### Estimating the interaction parameter under G-E independence

We evaluated bias, variance estimation, and confidence interval coverage for the interaction parameter for the three approaches under *G-E* independence. In these simulations, *S_D_* and *E* were independently normally distributed both with mean zero and variance 0.25. We simulated *D* as a Bernoulli random variable with success probability exp(*η*_0_ + 0.589*S_D_* − 0.5*E* − 0.1*S_D_E*), where *η*_0_ was chosen to yield a disease prevalence of 1% in the population. The value of the *S_D_* main effect parameter, 0.589, corresponds approximately to a PRS with modest predictive power.

The model fit under the case-only method was the model defined in equation 3 and the models fit under the case-control and cohort approaches were

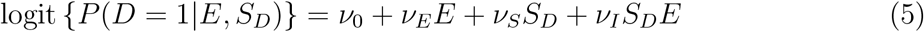

and

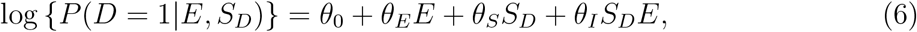

respectively. For the case-only method, the residual standard error estimate from the linear regression model was used to estimate *σ*^2^.

We also investigated the type I error rate of the three approaches. The simulation set-up was identical to that described above, except *θ_I_* = 0; that is, *D* was simulated as a Bernoulli random variable with success probability exp(*η*_0_ + 0.589*S_D_* − 0.5*E*), where again *η*_0_ was such that the disease prevalence in the population was 1%.

#### Estimating the main effect of S_D_ under G-E independence

We evaluated the estimates of the main effect of *S_D_* in terms of bias, variance estimation, and confidence interval coverage.

The simulation set-up was identical to that described above. For the case-only method, 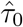 was centered by an estimate of *μ* from an external sample of size *n*\* (*n*\* = 5000,10000, or 50000). This centered estimate was then scaled by 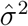, where 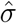 was the residual standard error estimate from the case-only linear regression model, yielding 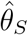. The standard error of 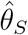 was estimated as described in Web Appendix D. We also evaluated the type I error rate of the three approaches by setting *θ_S_* to zero.

#### Estimating the interaction parameter under conditional G-E independence

We considered the setting where *S_D_* and *E* were independent conditional on *S_E_*. In particular, *S_D_* and *S_E_* were jointly normally distributed with mean zero, variances of 0.25 and 0.1, respectively, and covariance 0.05, yielding modest correlation between *S_D_* and *S_E_*. Then *E* was defined as 0.5*S_E_* + *ξ*, where *ξ* ~ *N*(0, 0.22); this means *S_E_* accounted for approximately 10% of the variation in *E* and *E* had a variance of approximately 0.25. The outcome *D* was simulated as a Bernoulli random variable with success probability exp(*η*_0_ + 0.589*S_D_* − 0.5*E* − 0.1*S_D_E*), where *η*_0_ was chosen to yield a disease prevalence in the population of 1%. For the case-only, case-control, and cohort approaches, we fit the regression models given in equations 4, 5, and 6, respectively. For the case-only method, the residual standard error estimate from the linear regression model was used to estimate *ψ*^2^. Finally, we investigated the type I error rate of the three approaches (i.e., we set *θ_I_* = 0).

## RESULTS

### Application to data from the UK Biobank

Although no significant interactions were identified, the results of the case-only analysis generally mirror those of the cohort analysis and differ notably from those of the case-control analysis (Table 1). In particular, the confidence intervals from the case-only and cohort analyses are similar and are 30-40% narrower than those from the case-control analysis. These results indicate the case-only approach can yield efficiency gains over the case-control approach and essentially recover the efficiency of a cohort study. These results are also in line with the earlier analysis by Rudolph et al. (24): while they found significant interactions between a categorical PRS and both height and alcohol intake, they did not observe a dose-response relationship, consistent with our findings of no significant interactions with a continuous PRS.

**Table 1.** Analysis of Interactions Between a 151-SNP Breast Cancer PRS and Environmental Variables in the UK Biobank.

| Environmental Variable | Analysis | Interaction Estimate<sup>a</sup> | 95% Confidence Interval |
| --- | --- | --- | --- |
| Alcohol<sup>b</sup> | Case-only | 0.998 | 0.960, 1.038 |
|  | Case-control | 1.024 | 0.965, 1.086 |
|  | Cohort | 1.001 | 0.965, 1.038 |
| BMI<sup>c</sup> | Case-only | 1.002 | 0.994, 1.010 |
|  | Case-control | 1.000 | 0.988, 1.011 |
|  | Cohort | 1.002 | 0.995, 1.009 |
| Height<sup>d</sup> | Case-only | 0.977 | 0.948, 1.008 |
|  | Case-control | 0.975 | 0.930, 1.023 |
|  | Cohort | 0.974 | 0.942, 1.007 |

Abbreviations: BMI, body mass index; PRS, polygenic risk score; SNP, single nucleotide polymorphism.
<sup>a</sup> For the case-only and cohort analyses, this is an estimate of the interaction on the relative risk scale. For the case-control analysis, this is an estimate of the interaction on the odds ratio scale. These parameters are expected to be approximately equal.
<sup>b</sup> Per 10 grams/day and standard deviation of the PRS (0.509).
<sup>c</sup> Per kilogram/meter<sup>2</sup> and standard deviation of the PRS (0.509).
<sup>d</sup> Per 5 centimeters and standard deviation of the PRS (0.509).

### Simulations

#### Estimating the interaction parameter under G-E independence

For all three approaches, the interaction parameter was estimated with little bias, the mean estimated standard errors were generally close to the empirical standard errors (i.e., the standard deviation of the parameter estimate across simulations), the coverage probabilities were approximately 95%, and the type I error rates were close to the nominal level (Table 2). The mean estimated standard errors for the case-only method were slightly lower than the empirical standard errors. These differences were small, diminished with increasing sample size, and were likely due to small sample bias associated with using an estimate of *σ*^2^ to obtain 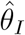. We also observe marked gains in efficiency (that is, reduced empirical variance) for the case-only method relative to the standard case-control analysis: examination of the variance ratios indicates the case-only method was more than twice as efficient as the case-control analysis and nearly recovered the efficiency of the cohort approach.

**Table 2.** Mean Parameter Estimate, Mean Estimated and Empirical Standard Errors, Coverage Probability of 95% Confidence Intervals, and Type I Error at the *α* = 0.05 Level for the Interaction Parameter for Data Simulated Under *G-E* Independence.

| $n$ | Analysis | Parameter Estimate<sup>a</sup><br>($\theta_I = -0.1$) | Standard Error | | Coverage<br>Probability | Type I<br>Error Rate |
| --- | --- | --- | --- | --- | --- | --- |
|  |  |  | Estimated | Empirical |  |  |
| 1000 | Case-only | -0.0996 | 0.1266 | 0.1314 | 0.939 | 0.054 |
|  | Case-control | -0.1040 | 0.1868 | 0.1879 | 0.950 | 0.051 |
|  | Cohort | -0.1018 | 0.1247 | 0.1257 | 0.948 | 0.048 |
| 2000 | Case-only | -0.1015 | 0.0896 | 0.0911 | 0.940 | 0.047 |
|  | Case-control | -0.1118 | 0.1319 | 0.1313 | 0.948 | 0.042 |
|  | Cohort | -0.0953 | 0.0886 | 0.0878 | 0.946 | 0.042 |
| 5000 | Case-only | -0.1006 | 0.0566 | 0.0571 | 0.948 | 0.045 |
|  | Case-control | -0.1076 | 0.0832 | 0.0864 | 0.947 | 0.047 |
|  | Cohort | -0.0981 | 0.0561 | 0.0548 | 0.950 | 0.071 |

Abbreviations: $E$ , environment; $G$ , gene.
<sup>a</sup> For the case-only and cohort analyses, this is an estimate of the interaction on the log relative risk scale. For the case-control analysis, this is an estimate of the interaction on the log odds ratio scale. These parameters are approximately equal in these simulations.

#### Estimating the main effect of S_D_ under G-E independence

All three methods did well in estimating the main effect of *S_D_* and its standard error and were able to maintain the nominal coverage probability and type I error rate (Table 3). Again, we see some discrepancies in the mean estimated and empirical standard error for the case-only method that lessen with increasing *n*. However, the mean estimated standard errors for the case-only method generally tracked well with the empirical standard errors, indicating we were able to adequately account for the use of an estimate of *μ*. The estimates from the case-only method were at least as efficient as those from the case-control approach and in many cases were substantially more efficient. In general, as *n*/n* grew, the efficiency of the case-only method relative to the case-control analysis increased and approached that of the cohort analysis. When *n*\* and *n* were of a similar magnitude, i.e., *n*\* = 5000, the estimates from the case-only method were up to 40% more efficient than those from the case-control approach (when *n*\* = *n* = 5000, the efficiency of the estimates from the case-only and case-control approaches was comparable). When *n*\* = 10, 000, that is, at least twice as large as *n*, the case-only method was between 28% and 45% more efficient. In the setting where *n*\* was at least ten times as large as *n* (*n*\* = 50, 000), the case-only method was between 62% and 81% more efficient.

**Table 3.** Mean Parameter Estimate, Mean Estimated and Empirical Standard Errors, Coverage Probability of 95% Confidence Intervals, and Type I Error at the *α* = 0.05 Level for the Main Effect of *S_D_* for Data Simulated Under *G-E* Independence.

| $n$ | Analysis | $n^*$ | Parameter Estimate<sup>a</sup><br>($\theta_S = 0.589$) | Standard Error | | Coverage<br>Probability | Type I<br>Error Rate |
| --- | --- | --- | --- | --- | --- | --- | --- |
|  |  |  |  | Estimated | Empirical |  |  |
| 1000 | Case-only | 5000 | 0.5879 | 0.0712 | 0.0765 | 0.932 | 0.049 |
|  |  | 10000 | 0.5891 | 0.0683 | 0.0758 | 0.924 | 0.047 |
|  |  | 50000 | 0.5894 | 0.0661 | 0.0679 | 0.936 | 0.059 |
|  | Case-control |  | 0.5945 | 0.0932 | 0.0914 | 0.957 | 0.056 |
|  | Cohort |  | 0.5880 | 0.0650 | 0.0673 | 0.944 | 0.043 |
| 2000 | Case-only | 5000 | 0.5903 | 0.0541 | 0.0564 | 0.941 | 0.051 |
|  |  | 10000 | 0.5885 | 0.0504 | 0.0556 | 0.919 | 0.056 |
|  |  | 50000 | 0.5906 | 0.0471 | 0.0499 | 0.931 | 0.048 |
|  | Case-control |  | 0.5968 | 0.0658 | 0.0635 | 0.956 | 0.051 |
|  | Cohort |  | 0.5881 | 0.0459 | 0.0474 | 0.946 | 0.062 |
| 5000 | Case-only | 5000 | 0.5890 | 0.0407 | 0.0417 | 0.942 | 0.049 |
|  |  | 10000 | 0.5887 | 0.0354 | 0.0369 | 0.945 | 0.044 |
|  |  | 50000 | 0.5896 | 0.0305 | 0.0321 | 0.930 | 0.048 |
|  | Case-control |  | 0.5968 | 0.0416 | 0.0418 | 0.939 | 0.049 |
|  | Cohort |  | 0.5887 | 0.0290 | 0.0286 | 0.951 | 0.052 |

Abbreviations: $E$ , environment; $G$ , gene.
<sup>a</sup> For the case-only and cohort analyses, this is an estimate of the main effect of $S_D$ on the log relative risk scale. For the case-control analysis, this is an estimate of the main effect of $S_D$ on the log odds ratio scale. These parameters are approximately equal in these simulations.

#### Estimating the interaction parameter under conditional G-E independence

When *S_D_* and *E* were independent conditional on *S_E_*, there was little bias, the mean estimated standard errors were generally similar to the empirical standard errors, the coverage probabilities were near 95%, and the type I error rates were close to 0.05 for all three approaches (Table 4). Again, we see large gains in efficiency for the case-only method relative to the case-control approach; in particular, the case-only method was at least 70% more efficient than the case-control approach and recovered much of the efficiency of the cohort approach.

**Table 4.** Mean Parameter Estimate, Mean Estimated and Empirical Standard Errors, Coverage Probability of 95% Confidence Intervals, and Type I Error at the *α* = 0.05 Level for Interaction Parameter for Data Simulated Under Conditional *G-E* Independence.

| $n$ | Analysis | Parameter Estimate<sup>a</sup><br>($\theta_I = -0.1$) | Standard Error | | Coverage<br>Probability | Type I<br>Error Rate |
| --- | --- | --- | --- | --- | --- | --- |
|  |  |  | Estimated | Empirical |  |  |
| 1000 | Case-only | -0.0992 | 0.1416 | 0.1405 | 0.956 | 0.046 |
|  | Case-control | -0.1077 | 0.1867 | 0.1883 | 0.955 | 0.052 |
|  | Cohort | -0.0998 | 0.1258 | 0.1240 | 0.952 | 0.057 |
| 2000 | Case-only | -0.1036 | 0.1001 | 0.1023 | 0.945 | 0.064 |
|  | Case-control | -0.1113 | 0.1317 | 0.1331 | 0.944 | 0.050 |
|  | Cohort | -0.1051 | 0.0893 | 0.0945 | 0.938 | 0.051 |
| 5000 | Case-only | -0.1021 | 0.0632 | 0.0618 | 0.950 | 0.049 |
|  | Case-control | -0.1048 | 0.0831 | 0.0811 | 0.955 | 0.057 |
|  | Cohort | -0.1026 | 0.0566 | 0.0542 | 0.961 | 0.045 |

Abbreviations: $E$ , environment; $G$ , gene.
<sup>a</sup> For the case-only and cohort analyses, this is an estimate of the interaction on the log relative risk scale. For the case-control analysis, this is an estimate of the interaction on the log odds ratio scale. These parameters are approximately equal in these simulations.

## DISCUSSION

We have proposed a method for case-only analysis of *G-E* interactions using a PRS. Our method is easy to implement, requiring only the ability to run linear regression. Importantly, the case-only approach incorporates an assumption of independence of the PRS *S_D_* and *E* (either marginally or conditionally given *S_E_*, a PRS for *E*), yielding marked gains in efficiency over logistic regression analysis of case-control data. The proposed method makes no assumptions about the distribution of *E*, lending flexibility to this approach. We used breast cancer data from the UK Biobank to illustrate our method, providing evidence it can offer gains in efficiency over the case-control analysis and produce results very similar to the analysis of the full cohort.

For most rare diseases, implementing a cohort study is not feasible; hence, a case-control study is frequently pursued. Our approach requires collecting only a sample of cases, yet it can nearly recover the results that would be observed in a cohort study. In addition to the gains in efficiency described above, the reliance on only a sample of cases is a strength of our method, as control selection issues may lead to bias in case-control studies (32–35). Additionally, if an estimate of the mean of *S_D_* in the underlying population is available (and, in the case of conditional independence, an estimate of the mean of *S_E_*), the main effect of *S_D_* can be estimated by our method, providing a fuller understanding of disease risk.

There are some similarities between our approach and recent work by Aschard et al. on methods for single-variant *G-E* interactions where E is continuous and measured with error (36). They demonstrated that a linear regression of *E* on *D, G*, and *DG* can be used to efficiently estimate the interaction parameter in this setting. However, their method does not leverage *G-E* independence. Furthermore, for evaluating *G-E* interaction with a PRS, it is more natural to model the PRS as the outcome as it is a continuous variable, while *E* could be categorical, ordinal, or continuous and may also be multidimensional.

Our method relies on an assumption of *G-E* independence to provide estimates of the interaction parameter *θ_I_* using only a sample of cases. We have shown how this assumption can be relaxed to allow for dependence between *S_D_* and *E* due to associations between the SNPs in *S_D_* and *E. S_D_* and *E* may also be correlated due to population stratification, as the distributions of both *E* and *S_D_* may vary by population strata (37,38). Since population stratification can also influence disease risk and possibly the association between *S_D_* and *D*, additional work is needed to develop efficient methods for studying interactions between *S_D_* and *E* in the presence of population stratification.

## ACKNOWLEDGEMENTS

Research reported in this article was funded through a Patient-Centered Outcomes Research Institute (PCORI) Award (ME-1602-34530). The statements, opinions in this article are solely the responsibility of the authors and do not necessarily represent the views of the Patient-Centered Outcomes Research Institute (PCORI), its Board of Governors or Methodology Committee.

The authors thank Haoyu Zhang for assistance in obtaining data from the UK Biobank.

Conflict of interest: none declared.

## Web Appendix A

### Derivation of conditional distribution of S_D_ in cases

Given (*S_D_|E*) ~ *S_D_* ~ *N*(*μ,σ*^2^), the distribution of the PRS in cases given *E* can be written

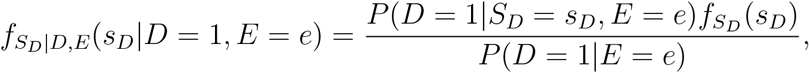

where *f* denotes a probability density. Under the model in equation 1, 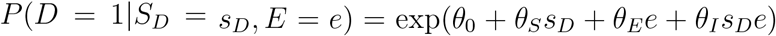 and 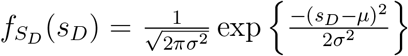. Furthermore, we can write

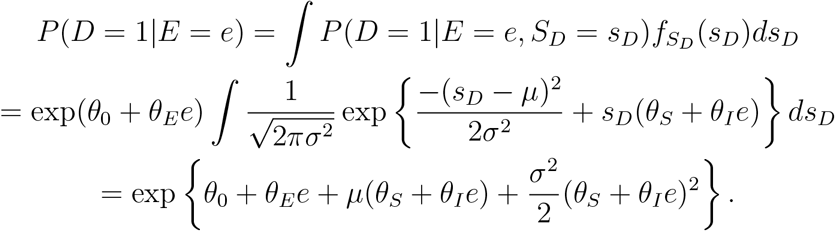

Then

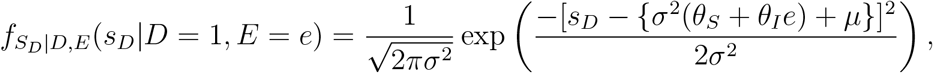

giving (*S_D_|E, D* =1) ~ *N*(*μ* + *σ*^2^(*θ_S_* + *θ_I_E*), *σ*^2^).

## Web Appendix B

### Derivation of variance of 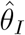 in case-only model

Consider the model presented in equation 3, *S_D_* = *τ*_0_ + *τ*_1_*E* + *ϵ*, where *τ*_1_ = *σ*^2^*θ_I_*. We propose using 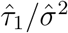 as an estimate of *θ_I_* and we claim that

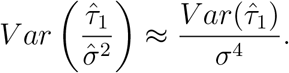

We can show this as follows. Using a first-order Taylor series approximation, we have

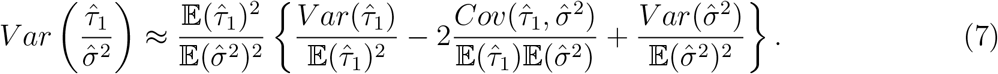

Consider the third term, 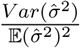. Recall that due to the central limit theorem, *S_D_* and (*S_D_|D* = 1, *E*) are approximately normally distributed. Thus, when *σ*^2^ is estimated from an external sample or a sample of controls or the residual standard error is used,

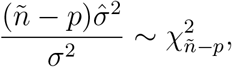

holds approximately, where *p* = 1 when an external sample or a sample of controls is used (and *ñ* is the size of that sample) and *p* =2 when the residual standard error from model in equation 3 is used (and *ñ* is the number of cases). This gives

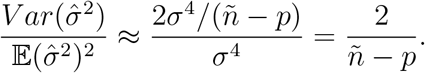

Next consider the second term, 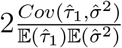. When *σ*^2^ is estimated in an independent sample (i.e., an external sample or an internal sample of controls), 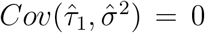. We claim that when 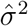 is the residual standard error from model in equation 3, 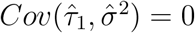. The score equations for (*τ*_0_, *τ*_1_, *σ*^2^) can be written as

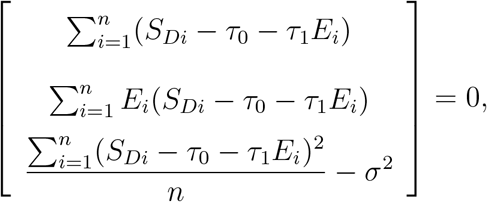

where *n* is the number of cases. The information matrix is then

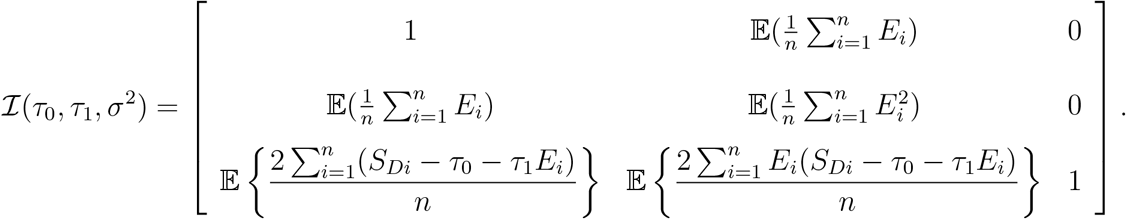

However,

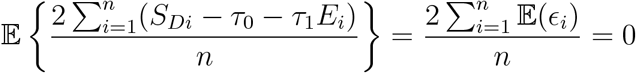

since the residuals have expectation zero. Likewise,

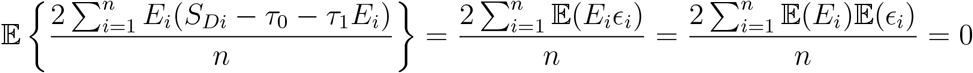

by independence of the residuals and *E* and the fact that the residuals have expectation zero. Thus,

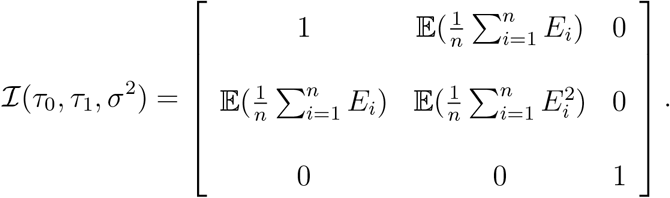

This gives 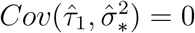, where 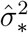 is the solution to the score equation, that is,

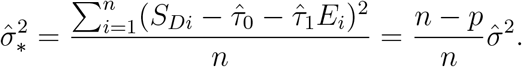

Then 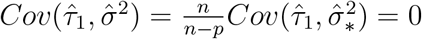, as claimed. This means that the second term in (7) is zero.

We can rewrite (7) as

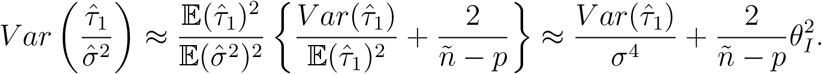

Provided |*θ_I_*| is not large (which is expected in general), we have

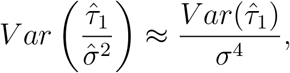

as claimed. We can then write

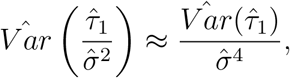

which can be used to obtain an estimate of the standard error of 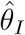.

## Web Appendix C

### Derivation of logit-risk model form under log-risk population model

The derivation in Web Appendix A is based on the log-risk population model. Suppose that a logit-risk model is fit to data generated by the log-risk population model. The population log-risk model can be written as

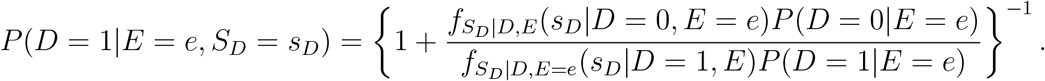

Then

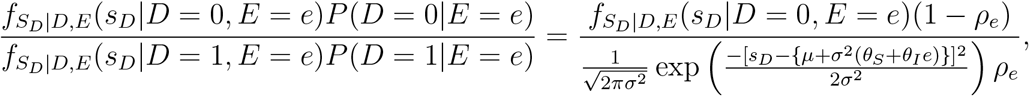

where 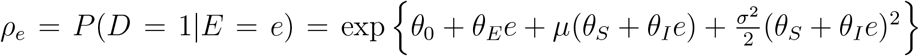. If the outcome is rare for all levels of *E, P*(*D* = 1|*E* = *e*) = *ρ_e_* ≈ 0 ∀ *e*, then *f_S_D_|D,E_*(*s_D_|D* = 0, *E* = *e*) ≈ *f_S_D_|E_*(*s_D_|E* = *e*) = *f_S_D__*(*s_D_*). This gives

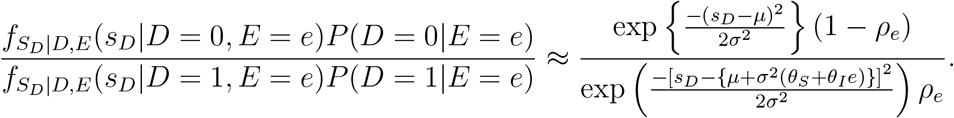

Then

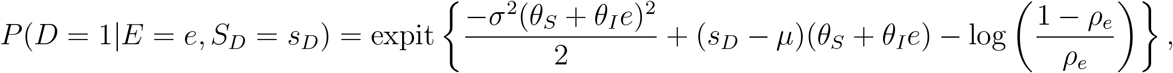

where 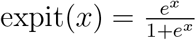, the inverse of the logit function. We can write

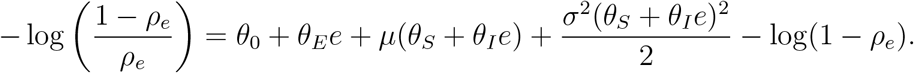

Combining this with the above, we have

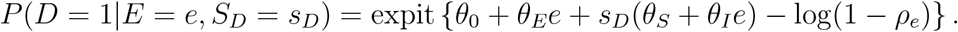

If the outcome is rare for each level of *E*, i.e., *ρ_e_* ≈ 0, then log(1 − *ρ_e_*) ≈ 0 ∀*e*, so the linear logit-risk model holds and the interaction on the relative risk scale is equivalent to the interaction on the odds ratio scale. When this does not hold, however, the two interaction parameters represent different population-level quantities.

## Web Appendix D

### Derivation of variance of 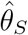 in case-only model when an estimate of μ is used

If *μ* is estimated using a sample of individuals, the estimated standard error of the PRS main effect must be adjusted to account for the estimation of *μ*. Without loss of generality, suppose 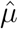 is used to center 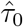: since *σ*^2^*θ_S_* = *τ*_0_ − *μ, θ_S_* can be estimated by 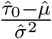. Then

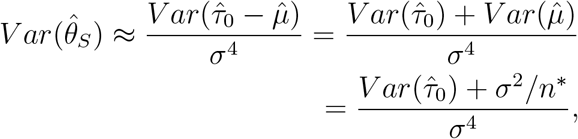

where the approximation follows from arguments similar to those in Web Appendix B, the first equality follows from the fact that *τ*_0_ and *μ* are estimated in independent samples (in a sample of cases and in either an external sample or an internal sample of controls, respectively) and so are uncorrelated, and *n*\* is the size of the sample used to estimate *μ*. Thus

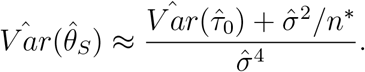

## Abbreviations

BMI: body mass index
*E*: environment
*G*: gene
GWAS: genome-wide association study
PRS: polygenic risk score
SNP: single-nucleotide polymorphism

## References

1. Yang Q and Khoury MJ. Evolving methods in genetic epidemiology. III. Gene-environment interaction in epidemiologic research. Epidemiol Rev. 1997;19(1):33–43.

2. Murcray CE, Lewinger JP, and Gauderman WJ. Gene-environment interaction in genome-wide association studies. Am J Epidemiol. 2009;169(2):219–226.

3. McAllister K, Mechanic LE, Amos C, et al. Current challenges and new opportunities for gene-environment interaction studies of complex diseases. Am J Epidemiol. 2017;186(7):753–761.

4. Khoury MJ. Emergence of gene-environment interaction analysis in epidemiologic research. Am J Epidemiol. 2017;186(7):751–752.

5. Ritz BR, Chatterjee N, Garcia-Closas M, et al. Lessons learned from past gene-environment interaction successes. Am J Epidemiol. 2017;186(7):778–786.

6. Duncan LE and Keller MC. A critical review of the first 10 years of candidate gene-by-environment interaction research in psychiatry. Am J Psychiatry. 2011;168(10):1041–1049.

7. London SJ and Romieu I. Gene by environment interaction in asthma. Annu Rev Public Health. 2009;30:55–80.

8. Simonds NI, Ghazarian AA, Pimentel CB, et al. Review of the gene-environment interaction literature in cancer: what do we know? Genet Epidemiol. 2016;40(5):356–365.

9. Carbone M, Amelio I, Affar EB, et al. Consensus report of the 8 and 9th Weinman Symposia on Gene x Environment Interaction in carcinogenesis: novel opportunities for precision medicine. Cell Death Differ. 2018;25(11):1885–1904.

10. Marley AR and Nan H. Epidemiology of colorectal cancer. Int J Mol Epidemiol Genet. 2016;7(3):105–114.

11. Rudolph A, Chang-Claude J, and Schmidt MK. Gene–environment interaction and risk of breast cancer. Br J Cancer. 2016;114(2):125–133.

12. Gauderman WJ, Mukherjee B, Aschard H, et al. Update on the state of the science for analytical methods for gene-environment interactions. Am J Epidemiol. 2017;186(7):762–770.

13. Piegorsch WW, Weinberg CR, and Taylor JA. Non-hierarchical logistic models and case-only designs for assessing susceptibility in population-based case-control studies. Stat Med. 1994;13(2):153–162.

14. Umbach DM and Weinberg CR. Designing and analysing case-control studies to exploit independence of genotype and exposure. Stat Med. 1997;16(15):1731–1743.

15. Chatterjee N and Carroll RJ. Semiparametric maximum likelihood estimation exploiting gene-environment independence in case-control studies. Biometrika. 2005;92(2):399–418.

16. Mukherjee B and Chatterjee N. Exploiting gene-environment independence for analysis of case-control studies: an empirical Bayes-type shrinkage estimator to trade-off between bias and efficiency. Biometrics. 2008;64(3):685–694.

17. Chatterjee N, Chen Y-H, Luo S, and Carroll RJ. Analysis of case-control association studies: SNPs, imputation and haplotypes. Stat Sci. 2009;24(4):489–502.

18. Gauderman WJ, Thomas DC, Murcray CE, et al. Efficient genome-wide association testing of gene-environment interaction in case-parent trios. Am J Epidemiol. 2010;172(1):116–122.

19. Chatterjee N, Shi J, and García-Closas M. Developing and evaluating polygenic risk prediction models for stratified disease prevention. Nat Rev Genet. 2016;17(7):392–406.

20. Kraft P and Aschard H. Finding the missing gene-environment interactions. Eur J Epidemiol. 2015;30(5):353–355.

21. Maas P, Barrdahl M, Joshi AD, et al. Breast cancer risk from modifiable and non-modifiable risk factors among white women in the United States. JAMA Oncol. 2016;2(10):1295–1302.

22. Garcia-Closas M, Gunsoy NB, and Chatterjee N. Combined associations of genetic and environmental risk factors: implications for prevention of breast cancer. J Natl Cancer Inst. 2014;106(11).

23. Garcia-Closas M, Rothman N, Figueroa JD, et al. Common genetic polymorphisms modify the effect of smoking on absolute risk of bladder cancer. Cancer Res. 2013;73(7):2211–2220.

24. Rudolph A, Song M, Brook MN, et al. Joint associations of a polygenic risk score and environmental risk factors for breast cancer in the Breast Cancer Association Consortium. Int J Epidemiol. 2018;47(2):526–536.

25. Yengo L, Sidorenko J, Kemper KE, et al. Meta-analysis of genome-wide association studies for height and body mass index in ~ 700,000 individuals of European ancestry. bioRxiv. 2018;(doi: https://doi.org/10.1101/274654). Accessed January 18, 2019.

26. Sudlow C, Gallacher J, Allen N, et al. UK Biobank: an open access resource for identifying the causes of a wide range of complex diseases of middle and old age. PLoS Med. 2015;12(3):e1001779.

27. UK Biobank. UK Biobank: Protocol for a large-scale prospective epidemiological resource. http://www.ukbiobank.ac.uk/wp-content/uploads/2011/11/UK-Biobank-Protocol.pdf. Accessed September 26, 2018.

28. Michailidou K, Lindstrom S, Dennis J, et al. Association analysis identifies 65 new breast cancer risk loci. Nature. 2017;551(7678):92–94.

29. Zou G. A modified poisson regression approach to prospective studies with binary data. Am J Epidemiol. 2004;159(7):702–706.

30. van den Brandt PA, Spiegelman D, Yaun SS, et al. Pooled analysis of prospective cohort studies on height, weight, and breast cancer risk. Am J Epidemiol. 2000;152(6):514–527.

31. Collaborative Group on Hormonal Factors in Breast Cancer. Alcohol, tobacco and breast cancer-collaborative reanalysis of individual data from 53 epidemiological studies, including 58515 women with breast cancer and 95067 women without the disease. Br J Cancer. 2002;87(11):1234–1245.

32. Wacholder S, McLaughlin JK, Silverman DT, et al. Selection of controls in case-control studies: I. Principles. Am J Epidemiol. 1992;135(9):1019–1028.

33. Wacholder S, Silverman DT, McLaughlin JK, et al. Selection of controls in case-control studies: II. Types of controls. Am J Epidemiol. 1992;135(9):1029–1041.

34. Wacholder S, Silverman DDT, McLaughlin JK, et al. Selection of controls in case-control studies: III. Design options. Am J Epidemiol. 1992;135(9):1042–1050.

35. Clayton D and McKeigue PM. Epidemiological methods for studying genes and environmental factors in complex diseases. Lancet. 2001;358(9290):1356–1360.

36. Aschard H, Spiegelman D, Laville V, et al. A test for gene-environment interaction in the presence of measurement error in the environmental variable. Genet Epidemiol. 2018;42(3):250–264.

37. Reisberg S, Iljasenko T, Lall K, et al. Comparing distributions of polygenic risk scores of type 2 diabetes and coronary heart disease within different populations. PLoS One. 2017;12(7):e0179238.

38. Martin AR, Gignoux CR, Walters RK, et al. Human demographic history impacts genetic risk prediction across diverse populations. Am J Hum Genet. 2017;100(4):635–649.

